# Heat stress does not induce wasting symptoms in the giant California sea cucumber (*Apostichopus californicus*)

**DOI:** 10.1101/2022.06.02.494586

**Authors:** Declan Taylor, Jonathan James Farr, Em G Lim, Jenna Laurel Fleet, Sara Jayne Smith Wuitchik, Daniel Michael Wuitchik

## Abstract

Oceanic heat waves have significant impacts on disease dynamics in marine ecosystems. A severe sea cucumber wasting event occurred in Nanoose Bay, British Columbia, Canada, following an extreme heat wave, resulting in mass mortality of *Apostichopus californicus*. Here, we sought to determine if heat stress in isolation could trigger wasting symptoms in *A. californicus*. We exposed sea cucumbers to i) a simulated marine heat wave (22ºC), ii) an elevated temperature treatment (17ºC), or iii) control conditions (12ºC). We measured the presence of skin ulcers, mortality, posture maintenance, antipredator defences, spawning, and organ evisceration during the 79-hour thermal exposure, as well as 7-days post-exposure. Both the 22°C and 17°C treatments elicited stress responses where individuals exhibited a reduced ability to maintain posture and an increase in stress spawning. The 22ºC heat wave was particularly stressful, as it was the only treatment where mortality was observed. However, none of the treatments induced wasting symptoms as observed in the Nanoose Bay event. This study provides evidence that sea cucumber wasting is not triggered by heat stress in isolation, leaving the cause of the mass mortality event observed in Nanoose unknown.

## Introduction

Climate change is increasing the intensity, duration, range, and frequency of marine heat waves across the globe with potentially catastrophic effects on organism fitness, ecosystems, and human economies (Frölicher, Fischer & Gruber, 2018; Allan et al., 2021). In marine ecosystems these extreme climatic events often cause immediate and mass mortality at all trophic levels from thermal stress, starvation, toxicity, and hypoxia (Cavole et al., 2016; Di Lorenzo & Mantua, 2016; von Biela et al., 2019; Suryan et al., 2021). Marine heat waves may exacerbate disease because thermal stress can compromise the immune response, and warmer temperatures can increase the virulence of pathogens (Harvell et al., 1999; Marcogliese, 2008; Branco et al., 2012; Matozzo et al., 2012; Oliver et al., 2017). This temperature-associated increase in virulence and infectivity has been associated with stimulating and intensifying sea star wasting disease to devastating effects (Harvell et al., 1999; Bates, Hilton & Harley, 2009; Eisenlord et al., 2016; Hewson et al., 2018; Aquino et al., 2021). A notorious example of the impacts of a wasting disease outbreak was the 2013-current sea star wasting epidemic, which has affected more than 20 sea star species in the Northeast Pacific Ocean over the last decade (Hewson et al., 2018). Sea star wasting disease encompases a broad set of symptoms including twisted arms, lesions, deflation/loss of turgor, loss of arms, lack of grip strength in tube feet, and liquefaction (Bates, Hilton & Harley, 2009; Menge et al., 2016; Hewson et al., 2018). In most sea star species, the driving cause of wasting disease remains largely uncertain. It has often been assumed that wasting in echinoderms is driven by infectious agents (Hewson et al., 2014; Bucci et al., 2017; Miner et al., 2018; Hewson, Johnson & Tibbetts, 2020). Warm temperatures have also been linked to several mass mortality events (Bates, Hilton & Harley, 2009; Eisenlord et al., 2016; Harvell et al., 2019). Because of the population-level impacts on ecologically important species, understanding the pathogenic and environmental drivers of sea star wasting remains an area of active research (Aalto et al., 2020; Aquino et al., 2021; Hewson, 2021).

Wasting is not limited to sea stars and is an emerging concern across closely related taxa. For example, sea urchins have faced a variety of disease-linked mass mortality events and epizootics (Feehan & Scheibling, 2014). Several of these bacterial and amoebic diseases have been linked to warm water anomalies, from climate events or storms (Sweet, 2020). Red sea urchins (*Mesocentrotus franciscanus*) and purple sea urchins (*Strongylocentrotus purpuratus*), have suffered from “bald sea urchin disease” and “sea urchin wasting disease” along the Pacific coast of British Columbia (B.C.). Pathology of these diseases includes lesions to the body wall and a shortening or loss of spines, and both have been associated with mass mortality events (Feehan & Scheibling, 2014; Sweet, 2020). While these diseases are believed to have bacterial origins, similarly to that of sea star wasting, a single bacterial strain has not been identified as the cause (Sweet, 2020). Disease-mediated mass mortality events in urchins and sea stars have caused trophic cascades and dramatic ecosystem shifts, highlighting the importance of understanding these ecological phenomena. Recent evidence has emerged that wasting may occur in sea cucumbers as well, promoting further concerns about the impacts of marine diseases in shallow, near-shore ecosystems.

Since 2014, there have been reports of giant California sea cucumbers (*Apostichopus californicus*) with wasting symptoms similar to those of sea stars throughout their range in the northeast Pacific Ocean (Hewson, Johnson & Tibbetts, 2020). Symptoms of wasting in sea cucumbers (Echinodermata: Holothuria) include non-focal lesions and fissures across the body wall, epidermal tissue sloughing, and rapid liquefaction (Hewson, Johnson & Tibbetts, 2020; Fig. 1C). Following a series of heat waves from June 25 to July 07 2021 (Environment & Climate Change Canada, 2021; Ocean Networks Canada) a wasting event with very high mortality occurred in the Strait of Georgia (near Nanoose, B.C.), from late August to October 2021 (Fig. 1D; Lim, *pers comm*). At its peak, up to 94% of observed *A. californicus* at a single site showed evidence of wasting. On average, across all affected sites, 50% of observed *A. californicus* exhibited wasting symptoms. This highlights a potential link between thermal stress and wasting in *A. californicus*, especially given that healthy individuals were found below depths of 19 meters where the water was cooler (Lim, *pers comm*). The effect of temperature-related stressors on wasting in sea cucumbers has yet to be tested.

**Figure 1.**
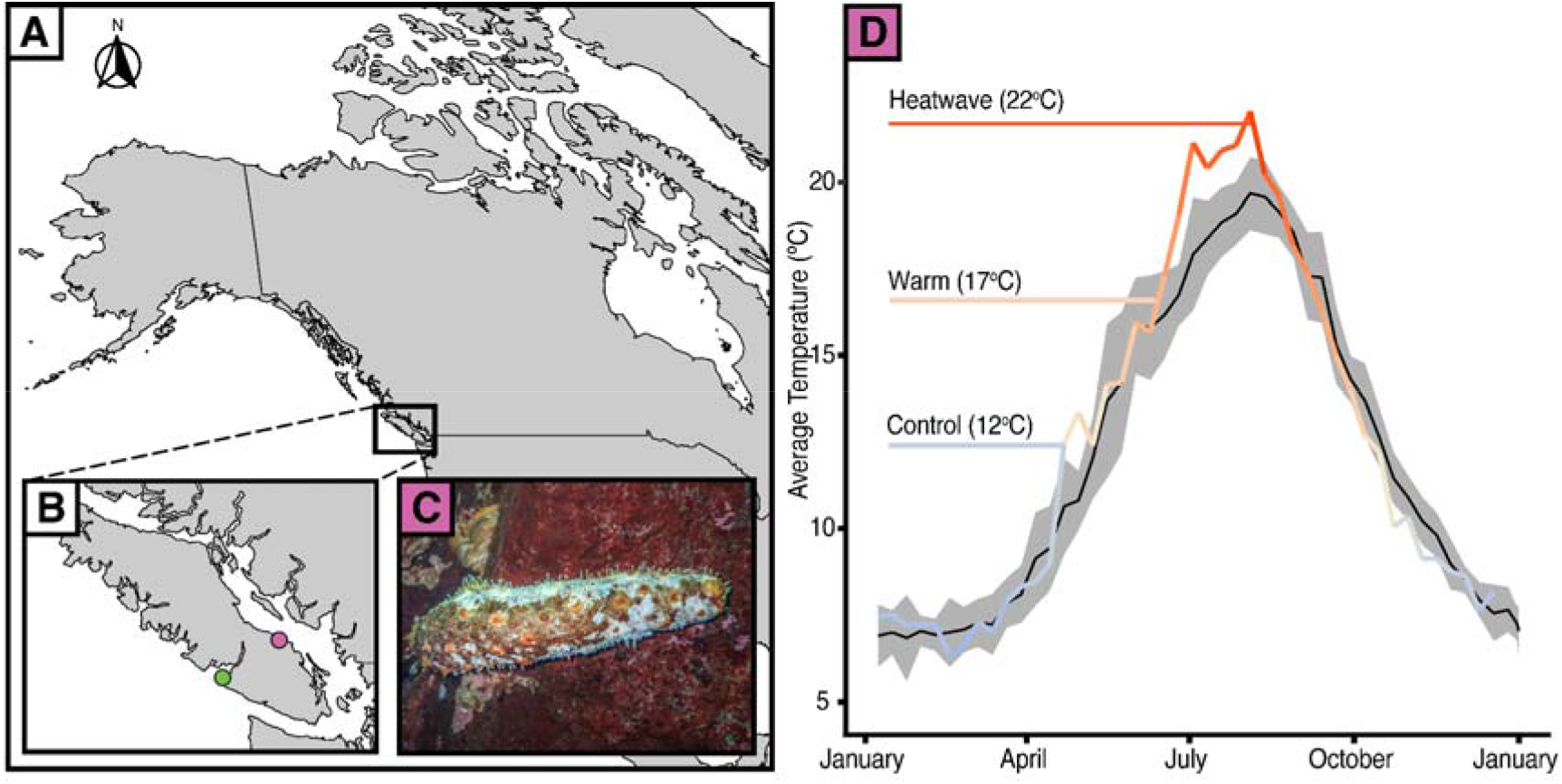
Temperature anomalies and sea cucumber wasting. A) Map of western North America with B) an inset of Vancouver Island showing the data collection locations of Bamfield Inlet (green), Nanoose Bay (purple). C) *A. californicus* in Nanoose Bay exhibiting wasting symptoms. D) 8-day rolling average seasonal sea surface temperatures in Nanoose Bay from 2010 to 2020 (black line with grey SE ribbon) and 2021 (colour gradient line), with experimental temperature treatments highlighted.

Here, we assess whether acute heat stress induces wasting in *A. californicus*. We also evaluate behavioural stress response signatures through stiffening, spawning, and evisceration to explore how they are affected by extreme elevated temperature. Sea cucumbers are important benthic detritivores that play an important role in organic matter decomposition and sediment aeration (Purcell, Conand & Byrne, 2016). Evidence of a connection between temperature and mass wasting events in sea stars (Bates, Hilton & Harley, 2009; Eisenlord et al., 2016; Harvell et al., 2019) necessitates a proactive assessment of whether heat stress can induce similar symptoms in sea cucumbers.

## Materials and Methods

### Collection and Acclimation

We collected *Apostichopus californicus* via SCUBA from a depth of 7-12 m in July 2021 from Scott’s Bay and Bamfield Inlet in Barkley Sound, British Columbia (48°50’02”N, 125°08’45”W). The sea cucumbers were initially collected and used in an unrelated short term tagging experiment at the end of July 2021, which had no measurable impact on sea cucumber behaviour or physiology (Leedham et al., unpublished data, Cieciel, Pyper & Eckert, 2009). The sea cucumbers were then maintained in an open flow-through system at the Bamfield Marine Sciences Centre, with inflow from Bamfield Inlet for four months prior to the start of the thermal stress experiment.

### Experimental Design

To determine an average wet weight of each individual, sea cucumbers were weighed and measured twice. We monitored if sea cucumbers were defecating throughout the experiment, as cessation of defecation is considered to be indicative of a loss or atrophy of digestive organs (Swan, 1961; Fankboner & Cameron, 1985). Individuals (*N* = 56) were randomly divided into one of three temperature treatments: control (12ºC, *n* = 19), warm (17ºC, *n* = 19), or heat wave (22ºC, *n* = 18). Our experimental temperatures reflect average summer temperatures and those recorded in the Strait of Georgia when wasting of *A. californicus* was observed (Fig. 1D). Specifically, the average temperature across July and August 2020 and 2021 was 17.5ºC at 5m depth and 11.9ºC at 20m depth (Fig. S1). In the summer 2021 heat wave, a maximum temperature at 5m depth of 21.6ºC was reached on August 04, 2021 (Pawlowicz, 2017; Xuereb et al., 2018; Chen et al., 2021; Ocean Networks Canada). Our control temperature (12ºC) was that of the Bamfield Inlet at the time of the temperature experiment and was the temperature the cucumbers were acclimated to.

We separated the sea cucumbers into 27 experimental containers (61 × 41 × 22.2 cm) with two individuals per container, aside from 2 smaller containers (33 × 45.7 × 11.4 cm) that housed one individual each. A plastic mesh divider was used to separate sea cucumbers within each container to allow for tracking individuals throughout the experiment. The containers were randomly placed into sea tables which acted as water baths allowing for different treatment temperatures. The target temperatures were achieved by having a constant flow of inlet sea water to the control sea tables (12ºC), ambient temperature water standing in the sea table (warm; 17ºC), or water heated with two 800W aquarium heaters (heat wave; 22ºC). The 22ºC treatment temperature was gradually increased over a 24-hour period in the sea table water baths (day 1; Fig. S1). The water remained at target temperatures for 79 hours, after which they were lowered back to the control temperature of 12ºC over 9-hours. Days 5 through 12 were a recovery period where all sea cucumber holding containers were maintained on the flow through system. Throughout the experiment, the sea cucumbers were closely monitored. Water changes were completed as needed to maintain nitrate and ammonium levels below 0.5 ppm, and fresh sea water was heated to the appropriate treatment temperature prior to water changes. Food was withheld during the thermal stress window.

### Phenotyping Sea Cucumber Stress

We assessed sea cucumbers for wasting symptoms throughout the experiment and counted skin ulcers on days 4, 5, 6, and 12. Ulcers were classified as either minor or major based on their size and appearance. We defined minor ulcers as small lesions on the ends of spines, without signs of considerable discoloration, and where the dermis was not fully damaged (Fig. 2B). We classified major ulcers as visibly open wounds exposing white tissue beneath (Fig. 2D).

**Figure 2.**
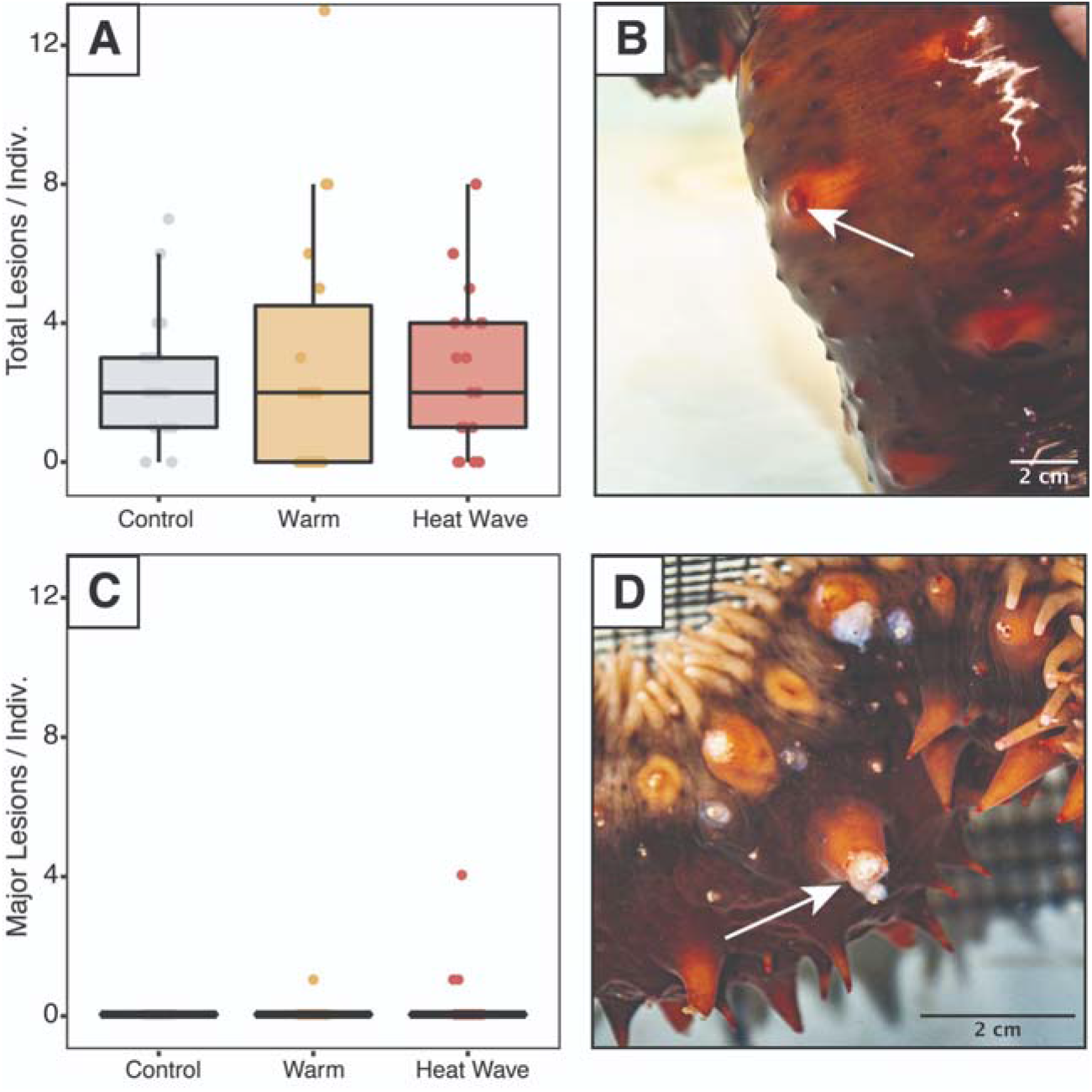
Skin ulcers observed in *Apostichopus californicus* across treatments. A) Minor ulcer count, defined as small lesions on the ends of spines (B).C) Major ulcer count, defined as open wounds that expose white subdermal tissue (D).

We assessed body stiffness using a 3-point ordinal scale to measure heat-stress induced deviations from normal behaviour. To do so, first we removed each individual from their bin and gently palpated them for 10 seconds to encourage them to stiffen. Then we placed them on an elevated platform to measure their ability to maintain their posture over 5 seconds. We assigned a score of 0 if the organism failed to stiffen at all (full droop), a score of 1 if it did not remain stiff for the full 5 seconds when placed on the platform (partial droop), and a score of 2 if it maintained its posture for the entire 5 seconds (no droop). Stiffness was measured on days 1-5 of the experiment (as a baseline and throughout the treatment), on day 7 (48 hours after experimental endpoint) and day 12 (7 days after experimental endpoint).

As some sea cucumbers spawned, we recorded whether there was any evidence of spawning every 12 hours. Spawning was assigned to each container rather than individual, since some individuals were co-housed (*N*_bin_ = 30). We also evaluated whether specimens had eviscerated based on the presence or absence of ejected internal organs during these routine checks (Fig. 4).

### Statistical Analyses

All statistical analyses were conducted in R (v4.0.3, R Core Team, 2020). We tested the distribution of the number of ulcers using the *fitDist* function (Rigby & Stasinopoulos, 2005) and determined that minor ulcer count best fit a geometric distribution. We modelled the maximum number of minor ulcers as a function of treatment, weight, evisceration (binary), and defecation status (binary). We included the sea table and bin as random effects. We then used backwards selection with the *stepGAIC* function from GAMLSS (Rigby & Stasinopoulos, 2005) to determine the most parsimonious combination of variables that best explained the maximum number of ulcers.

We examined the covariates that affected the likelihood of mortality with a logistic regression model. Sea cucumber mortality was a binary measure, with treatment, evisceration, defecation status, initial droop score, and initial weight as explanatory variables. Terms for the final model were selected via backwards model selection using *stepGAIC* (Rigby & Stasinopoulos, 2005) to produce the most parsimonious model of variation in sea cucumber mortality.

To assess changes in stiffness, we constructed full ordinal regression models using the *clmm* function (Christensen, 2019) with temperature treatment, date, and the interaction between treatment and date as explanatory variables. We restricted stiffness measurements to those taken before the heat treatment began (day 1), during the treatment (days 2-4) and immediately after the heat treatment (day 5). We included individual identity as a random effect to account for repeated measures on the same individuals over time. We also included bin and sea table as random effects to account for the paired (co-housing) and blocked (five bins per sea table) experimental design. We used backwards model selection to determine the most parsimonious models using *dredge* (Bartoń, 2020).

To assess the drivers of evisceration, we constructed a logistic regression model with treatment, weight, and defecation status as explanatory variables, along with sea table as a random effect. We determined the most parsimonious model through backwards selection. To compare the incidence of stress spawning across temperature treatments, we conducted a Dunn’s Kruskal-Wallace (K-W) test using the *dunnTest* function from the FSA package (Ogle et al., 2021).

All data and annotated code for the analyses described above are publicly available at https://github.com/declan-taylor/sea_cucumber_wasting.

## Results

The temperature treatments varied slightly from the target temperatures during the experimental heat waves. During the 79h treatment, the mean temperature of the control treatment was 12.4°C and varied from 10.8 - 14.0°C; the mean of the warm treatment was 16.6°C and ranged from 14.8°C to 17.9°C; the mean of the heat wave treatment was 21.7°C and varied from 19.6°C to 23.3°C. Temperature treatments were significantly different from each other (K-W χ^2^ = 463.32, df = 2, p < 2.2e^-16^).

Minor skin ulcers occurred during the experiment in all three treatments (*n*_*control*_ = 17, *n*_*warm*_ = 15, *n*_*heat wave*_ = 10; Fig. 2). Major ulcers were only observed in the warm (*n* = 1) and heat wave treatments (*n* = 2). These major ulcers appeared to heal throughout the recovery period and were re-classified as minor ulcers on day 12. The maximum number of minor ulcers per individual was not significantly explained by treatment, weight, evisceration, or defecation status.

Mortality was only observed in the 22°C heat wave treatment (Fig 3; *n* = 5), while there were no mortalities observed in the control or warm treatments. Based on backwards model selection, mortality was driven by treatment and weight (Table S1). Body stiffness was lower in the warm and heatwave treatments compared to the control treatment (Fig. 3). Backwards-selected models indicated that temperature treatment and day affected stiffness (Table S2). Sea cucumbers were significantly less likely to have high stiffness scores relative to the control treatment in the warm (p = 1.99e^-7^) and heat wave (p = 2.44e^-11^) treatments. Structural stiffness values were significantly likely to be lower than day 1 on day 3 (p = 1.37e^-5^), day 4 (p = 2.50e^-5^) and day 5 (p = 8.66e^-5^), but not on day 2 (p = 0.0627; Table S2).

**Figure 3.**
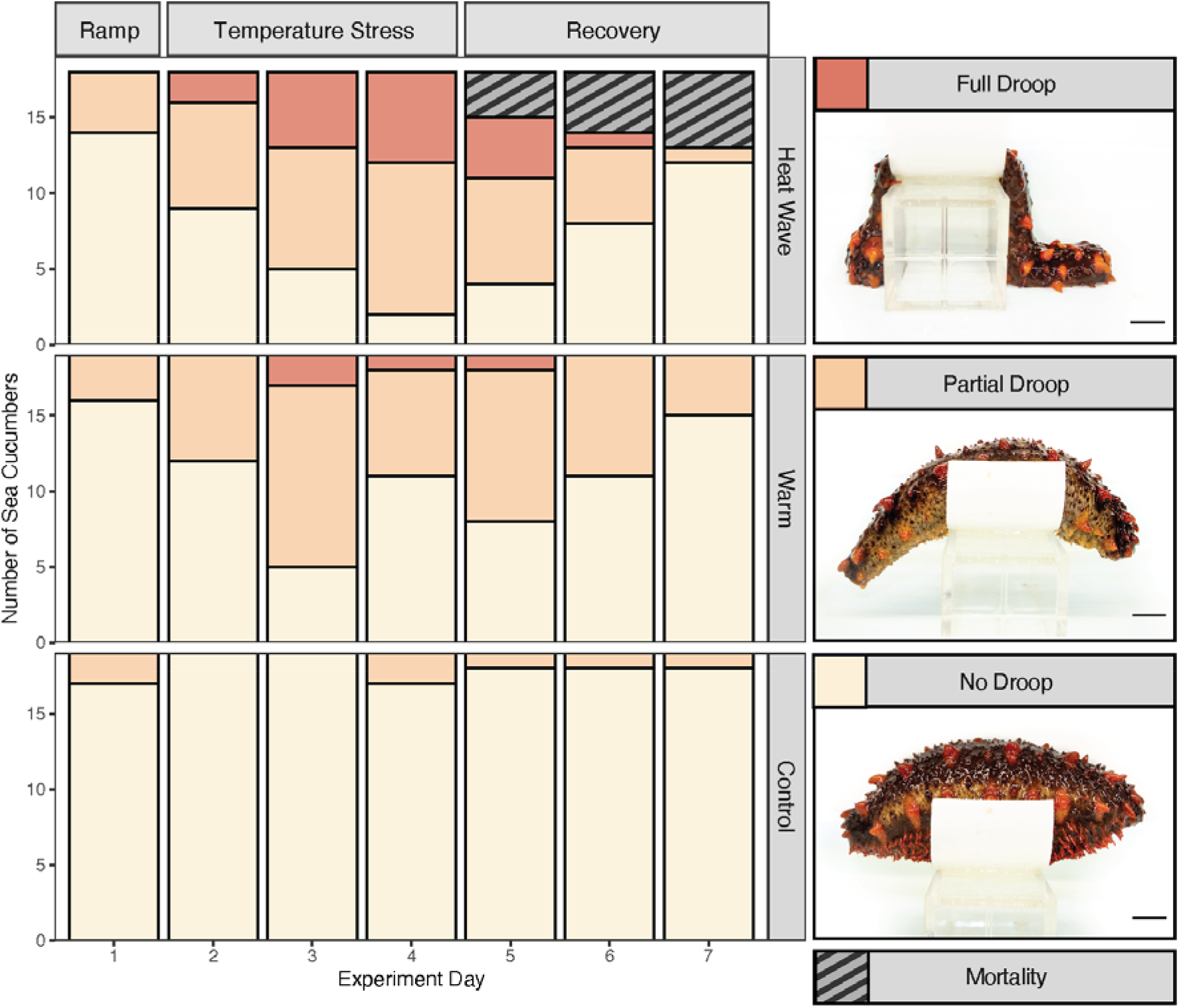
*Apostichopus californicus* stiffness and mortality at heat ramp (day 1), during temperature stress (days 2 - 4), and recovery (days 5 - 7) from the temperature treatment.

Stress spawning was observed during the temperature treatments (*n =* 11 bins), with most of the stress spawning occurring in the 17°C warm (*n =* 5 bins) and the 22°C heat wave (*n =* 4 bins) treatments. However, there was no significant difference in spawning between temperature treatments (K-W χ^2^ = 1.94, df = 2, p = 0.379).

Evisceration was observed across all treatments (*n*_*control*_ = 2, *n*_*warm*_ = 5, *n*_*heat wave*_ = 5). Treatment temperature did not explain a significant amount of the variance in the occurrence of evisceration (Table S3); however, weight (p = 0.0383) and defecation status (p = 0.0163) were significant drivers increasing likelihood of evisceration (Table S3). Along with evisceration of internal organs (Fig. 4A), we also observed the expulsion of the respiratory tree in two individuals (Fig. 4B). Both individuals were in the 22ºC heat wave treatment and mortality shortly followed the expulsion of the respiratory tree.

**Figure 4.**
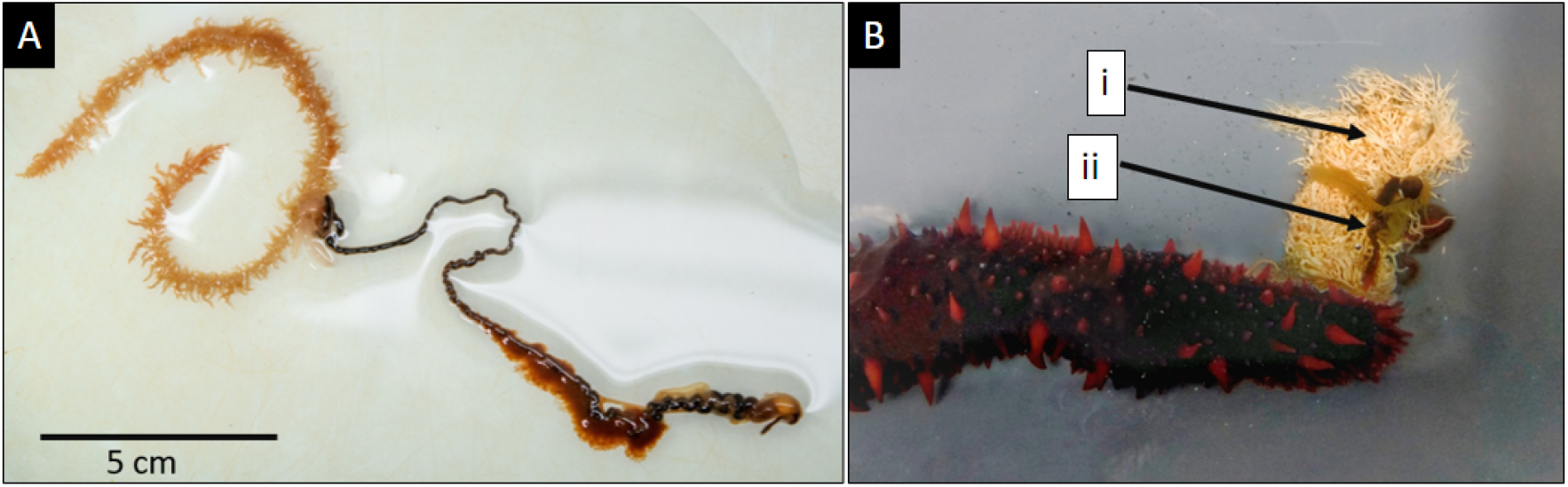
The eviscerated internal organs and respiratory tree of *Apostichopus californicus*. A) Eviscerated internal organs without respiratory tree expulsion compared to B) where both the i) respiratory tree and ii) internal organs were eviscerated.

## Discussion

The objective of our study was to determine if heat stress can induce wasting symptoms in *Apostichopus californicus*. While we saw minor skin ulcers at all treatment levels, and major ulcers in the warm and heat wave treatments, these are not characteristic wasting symptoms (Hewson et al., 2020; Fig. 1). Neither the minor or major ulcers that we observed matched the wasting symptoms reported in *A. californicus* in Nanoose Bay, B.C. (Lim, *pers comm*), or the isolated wasting events reported throughout the Pacific coast (Hewson, Johnson & Tibbetts, 2020). Both types of ulcers were very different from the white open lesions and fissures covering the dorsal body wall of wasting *A. californicus* in Nanoose (Fig. 1C). We also did not see any sloughing of body tissues or liquefaction, as has been anecdotally reported in previous wasting events (Schroeder, 2017; Hewson, Johnson & Tibbetts, 2020). Despite sharing some resemblance in colour, texture, and location to wasting disease symptoms, the sea cucumbers did not exhibit the full suite of symptoms that is typical from a wasting cucumber. Furthermore, unlike reports of widespread mortality resulting from wasting in wild *A. californicus*, the major ulcers in both of our specimens healed within the 7-day recovery period, and there was no evidence of these symptoms spreading to co-housed individuals. As such, there is no evidence that the sea cucumbers in our experiment were afflicted by the fatal wasting disease that has been previously reported (Schroeder, 2017; Hewson, Johnson & Tibbetts, 2020).

While we are uncertain of the ultimate cause of the ulcers that we observed, it is possible they were associated with handling. During the experiment, specimens were handled daily and measured for their posture maintenance, increasing the frequency of potential abrasions on the epidermal surface tissue. Major ulcers may have begun as minor ulcers that were then exacerbated by the high physiological stress caused by the 22°C treatment. White skin ulcerations, like the major ulcers we observed, are a recognized condition in other Holothuroidea, described as a Skin Ulceration Disease or Skin Ulceration Syndrome (SUS; Delroisse et al., 2020). SUS has been documented in commercially farmed *Apostichopus japonicus* and *Holothuria scabra*, and has been characterized by white ulcers on both sides of the body wall (Wang et al., 2007; Deng et al., 2009; Li et al., 2012; Zhang et al., 2018). Minor SUS symptoms in commercially farmed *A. japonicus* and the major ulcers in our *A. californicus* specimens are visually similar (Deng et al., 2009; Zhang et al., 2018). Unlike the SUS symptoms reported in *A. japonicus*, we did not see any indication of swelling or discolouration of the peristomes, and we did not see an initial abundance of ulcers around the mouth or cloaca (Becker et al., 2004; Wang et al., 2007; Delroisse et al., 2020). Extreme cases of SUS in farmed *A. japonicus* also bear resemblance to the wasting symptoms in wild *A. californicus*, raising further questions about the causes of skin ulceration in Holothuroidea (Delroisse et al., 2020).

Despite not seeing evidence of wasting, we saw strong evidence of whole organism and behavioural response to thermal stress. We observed five mortality events in the 22°C heat wave treatment. Though all mortalities occurred in the 22°C heat wave treatment, none of the mortalities were co-housed together, suggesting that the cause was unlikely to be a contagious disease or due to poor water quality.

Therefore, we suggest that the heat wave treatment is close to the upper critical thermal tolerance of *A. californicus*, but the warm treatment temperature does not cause sufficient harm for mortality to occur. Our findings align with previous work on larval life stages, where Ren et al. (2018) found that at 22°C, larval *A. californicus* experienced reduced survival which was not observed at 16°C or 18°C.

Beyond lethal effects, we observed reduced stiffening behaviour associated with both the 22°C heat wave and 17°C warm water treatment. A temperature-induced loss in stiffness may have implications for sea cucumber fitness under warming sea temperatures, as a limited ability to stiffen may inhibit their ability to avoid predation or maintain posture while feeding and spawning. Thermal stress may have reduced stiffness by causing muscular fatigue and relaxation (Dowd & Somero, 2013) in the circular and longitudinal-ambulacral muscles (Gao & Yang, 2015).

Stiffening is also caused by protein-mediated changes in mutable collagenous tissue within the dermis of sea cucumbers (Takehana et al., 2014). Therefore, heat stress may have reduced stiffening by denaturing or decreasing the production of tensilin, a stiffening protein, or increasing the production of the de-stiffening protein softenin (Yamada et al., 2010; Takehana et al., 2014; Tamori et al., 2016).

Unlike stiffening behaviour, we did not observe any significant trends in spawning behaviour or evisceration. Of the 11 spawning events, 9 occurred in elevated-temperature treatments (warm or heat wave) and this trend, although insignificant, was expected as stress spawning has been previously reported in other sea cucumber species (Battaglene et al., 2002; Rakaj et al., 2018; Schagerström et al., 2021). We were unable to associate spawning with individual phenotypes due to the co-housing experimental design. Evisceration appeared random across treatments, potentially because *A. californicus* in all treatments may have been reacting to handling stimulation and stress during the experiment (Ding et al., 2019). However, there is evidence that biological mechanisms, weight and defecation status, partially explain the non-treatment related variation in evisceration (Table S1). Defecation status showed that *A. californicus* that were not defecating were more likely to eviscerate. This may have occurred because the energetic cost of eviscerating digestive organs would be lower for *A. californicus* that had already ceased using their organs, either because they were preparing to eviscerate (Swan, 1961) or undergoing viscera atrophy (Fankboner & Cameron, 1985). As such, when stressed by handling, *A. californicus* that had already begun seasonal reductions in digestive function were more likely to eviscerate from stress and overstimulation. Unlike digestive tract evisceration, we do not believe that the expulsion of the respiratory tree in the 22°C heat wave treatment was linked to seasonal senescence (Fig. 4B). For both individuals, respiratory evisceration was followed by mortality, suggesting that this is an indication of extreme physiological distress from thermal exposure.

Since we observed skin ulcers under thermal stress that were not associated with wasting, other factors are likely causing recent wasting outbreaks in *A. californicus*. Like wasting, severe cases of SUS are highly transmissible and result in mortality (Delroisse et al., 2020); bacterial and viral pathogenic causal agents have been previously linked to SUS and wasting-like symptoms, both in other sea cucumbers (Deng et al., 2008, 2009; Liu et al., 2010; Zhang et al., 2018; Delroisse et al., 2020) and sea stars (Hewson et al., 2014, 2018; Work et al., 2021). A study examining a single wasting *A. californicus* specimen found a high viral load, but was unable to identify a specific pathogen causing wasting symptoms (Hewson et al., 2020).

Avenues for future research on wasting diseases in *A. californicus* could address the potential for shared pathology with SUS, given the symptomatic similarities, as well as the possibility of pathogenic origins. These investigations would add valuable insight to the field of wasting diseases, given the scarcity of published information on these events in echinoderms.

Studies conducted on historically asymptomatic populations that are isolated from wasting outbreaks could provide insight into whether the causal agents of wasting are naturally present in the *A. californicus* virome and/or microbiome. The specimens used here were from an asymptomatic *A. californicus* population that is genetically distinct from the Strait of Georgia populations. However, this population still receives substantial genetic influx (Xuereb et al., 2018), so we do not expect genetic diversity to confer differential vulnerability to wasting in our specimens compared to those in the Nanoose. Biotic factors (viral, bacterial) and abiotic factors (chemical pollution, hypoxia, eutrophication) should both be investigated because widespread environmental degradation and anthropogenic climate change are shifting pathogenic dynamics globally (Marcogliese, 2008; Allan et al., 2021).

In this study, we exposed *A. californicus* to extreme thermal stress as measured by mortality, degraded stiffening behaviour, and the development of skin ulcers. Despite this, we found no evidence that wasting is induced by temperature stress alone.

Therefore, the August 2021 mass wasting event in Nanoose, British Columbia, was likely not triggered solely by the anomalous heat wave. Determining the factors that cause and exacerbate wasting in *A. californicus* is essential for predicting and managing mass mortality events. Sea cucumbers are ecologically important benthic detritivores, which break down organic matter, recycle nutrients, and maintain sediment health (Wheeling, Verde & Nestler, 2007; Purcell, Conand & Byrne, 2016). Efforts to protect, manage, and sustainably harvest *A. californicus* in the face of global environmental change will require a comprehensive understanding of their stress responses, disease dynamics, and the novel threat of sea cucumber wasting.

## Acknowledgments

The species collections and experiments took place on the traditional territories of the Huu-ay-aht First Nations, a Nuu-chah-nulth Nation and signatory to the Maa-nulth First Nations Final Agreement, and we are grateful for the opportunity to conduct research in protected and sacred areas. We would like to thank the Bamfield Marine Sciences Centre for the resources and support required to conduct this research; Chloe Curry, Arya Horon, and Juliane Jones for their time providing lab assistance; Payton Arthur, Mike Chung, Gabrielle Languedoc, Sammie Foley, Juliane Jones, and Carter Burtlake for feedback on early versions of the manuscript.

## Supplementary Information

**Table S1.** Results from logistic regression model examining which variables best predicted sea cucumber mortality

| Variable | Coefficient | Std. Error | T | p |
| --- | --- | --- | --- | --- |
| (intercept) | -21.146 | 170.49 | -0.124 | 0.902 |
| Treatment: 17C | 0.0363 | 246.59 | 0 | 0.999 |
| Treatment: 22C | 14.72 | 170.42 | 0.086 | 0.932 |
| Weight | 0.0103 | 0.00553 | -1.86 | 0.0678 |

\*p < 0.05, \*\*p < 0.01

**Table S2.** Results of ordinal regression model examining the effect of treatment and experiment day on sea cucumber stiffness.□

| Variable | Coefficient | Std. Error | T | p |
| --- | --- | --- | --- | --- |
| Treatment: 17C | -2.99 | 0.575 | -5.20 | 1.99e-07** |
| Treatment: 22C | -4.05 | 0.606 | -6.68 | 2.44e-11** |
| Day 2 | -0.976 | 0.524 | -1.86 | 0.0627 |
| Day 3 | -2.30 | 0.528 | -4.35 | 1.37e-05** |
| Day 4 | -2.22 | 0.527 | -4.21 | 2.50e-05** |
| Day 5 | -2.12 | 0.541 | -3.93 | 8.66e-05** |

\*p < 0.05, \*\*p < 0.01

**Table S3.** Results from a backwards-selected logistic regression model examining the effect of evisceration as a function of defecation status and weight. Asterisks indicate significant effects.

|  | Coefficient | Std. Error | t-value | p |
| --- | --- | --- | --- | --- |
| (Intercept) | 1.713 | 1.128 | 1.52 | 0.135 |
| Defecating (yes) | -2.78 | 1.12 | -2.48 | 0.0163 * |
| Weight | -0.00431 | 0.00203 | -2.12 | 0.0383 * |

\*p < 0.05, \*\*p < 0.01

